# Simulating Scalp EEG from Ultrahigh-Density ECoG Data Illustrates Cortex to Scalp Projection Patterns

**DOI:** 10.1101/2025.06.24.660870

**Authors:** Seyed Yahya Shirazi, Julie Onton, Scott Makeig

**Affiliations:** Swartz Center for Computational Neuroscience, Institute for Neural Computation, University of California San Diego

## Abstract

Ultrahigh-density electrocorticography (μECoG) provides unprecedented spatial resolution for recording cortical electrical activity. This study uses simulated scalp projections from an μECoG recording to challenge the assumption that channel-level electroencephalography (EEG) reflects only local field potentials near the recording electrode. Using a 1024-electrode μECoG array placed on the primary motor cortex during finger movements, we applied Adaptive Mixture Independent Component Analysis (AMICA) to decompose activity into maximally independent grid activity components and projected these to 207 simulated EEG scalp electrode channels using a high-definition MR image-based electrical forward-problem head model. Our findings demonstrate how cortical surface-recorded potentials propagate to scalp electrodes both far from and near to the generating location. This work has significant implications for interpreting both EEG and ECoG data in clinical and research applications.

**Clinical Relevance:** This study provides insights for interpreting scalp EEG data, demonstrating that scalp channel activity represents a complex mixture of distributed cortical source activities rather than primarily activity generated nearest to the scalp electrodes. These findings may hopefully spur improvement in EEG-based diagnostics for neurological disorders.

## 1. Introduction

The local field potential (LFP) activity, dominated by population-level activity in the cortical neuropil, is typically considered to be the source of electrical activity recorded at the cortex and at scalp EEG electrodes. Through the last half century numerous studies have attributed (or assumed) channel-level EEG data to be generated by LFP activity ‘under’ the recorded EEG electrode(s), despite known volume conduction effects that blur EEG spatial resolution [1], [2]. This approach persists in the methods used in psychophysiological studies as well as clinical and research settings [3].

Source-level EEG analysis has revealed cortical and subcortical activities beyond the resolution of scalp-channel based analysis [Bathelt2014, Eom2022]. Basic biophysics demonstrates that every scalp EEG signal sums activities projected through volume conduction from many cortical areas [4], [5]. Techniques including Independent Component Analysis (ICA) can be used to separate source-area contributions arising near-independently in different cortical areas [6], [7].

This study investigates assumptions about using channel-level EEG to exclusively determine cortical activity near the recording area. We used ultra-high-density 1-mm spaced electrocorticography (μECoG) from a 32 × 32 mm patch on primary motor cortex [8] as ‘ground truth’ cortical electrical activity. We hypothesized that μECoG activity would reveal only a few “static” LFP sources appearing local to the recording area. Based on our previous computational simulations, we further hypothesized that this activity, projected through an electrical forward problem head-model, would be detected at EEG electrodes far beyond the immediate vicinity, illustrating how channel-level EEG only partially reflects cortical activity immediately underlying the channel electrodes.

## II. Methods

We analyzed a μECoG dataset reported by Tchoe et al. The data were collected using a custom platinum nanorods (PtNR) high-density electrode array with 1024 electrodes in a 32 × 32 mm patch with 1-mm inter-electrode spacing during an operation to plan surgery for epilepsy. The electrodes sat flush on the cortex without penetration. The patch included perfusion holes for cerebrospinal fluid passage. Initial recording was at 20 kHz using Intan RHD. We specifically analyzed mid-operational data from recording during a finger-tapping experiment lasting 9.6 minutes. ECoG and EEG processing analysis were performed using MATLAB (Mathworks Inc., Natick, MA) and the EEGLAB environment [9]. μECoG scalp projection and analysis were performed using Python (v3.10) and the Visualization Toolkit (vtk, v 9.0.3) [10].

### A. Preprocessing

The data were first downsampled to 1000 Hz and high-pass filtered at 0.5 Hz to remove DC offset. The variance across electrode banks was normalized to minimize imposed electrical effects. Unusable electrodes were identified and removed using visual inspection, correlation to any other channel <0.1, and variance above 7000 (empirical thresholds), retaining 807 of the 1024. Then Adaptive Mixture Independent Component Analysis (AMICA) [Palmer 2007] was applied to decompose μECoG activity into temporally independent components (‘grid ICs’). AMICA has demonstrated superior performance in identifying independent sources of EEG data whose scalp projection patterns (scalp maps) resemble single equivalent dipole projections compatible with generation in a single, likely compact cortical area [7].

### B. μECoG ICA analysis

Grid IC maps and activations were evaluated for recording noise patterns. Maps of ‘Cortical IC’ activity were characterized by smooth spatial gradients and relatively large (often 1-2 cm in diameter), irregularly shaped activity areas, while ‘artifact-IC’ maps included isolated electrode channels that were highly-weighted compared to signals at neighboring (within 2 mm) grid locations (see Fig. 1).

**Figure 1:**
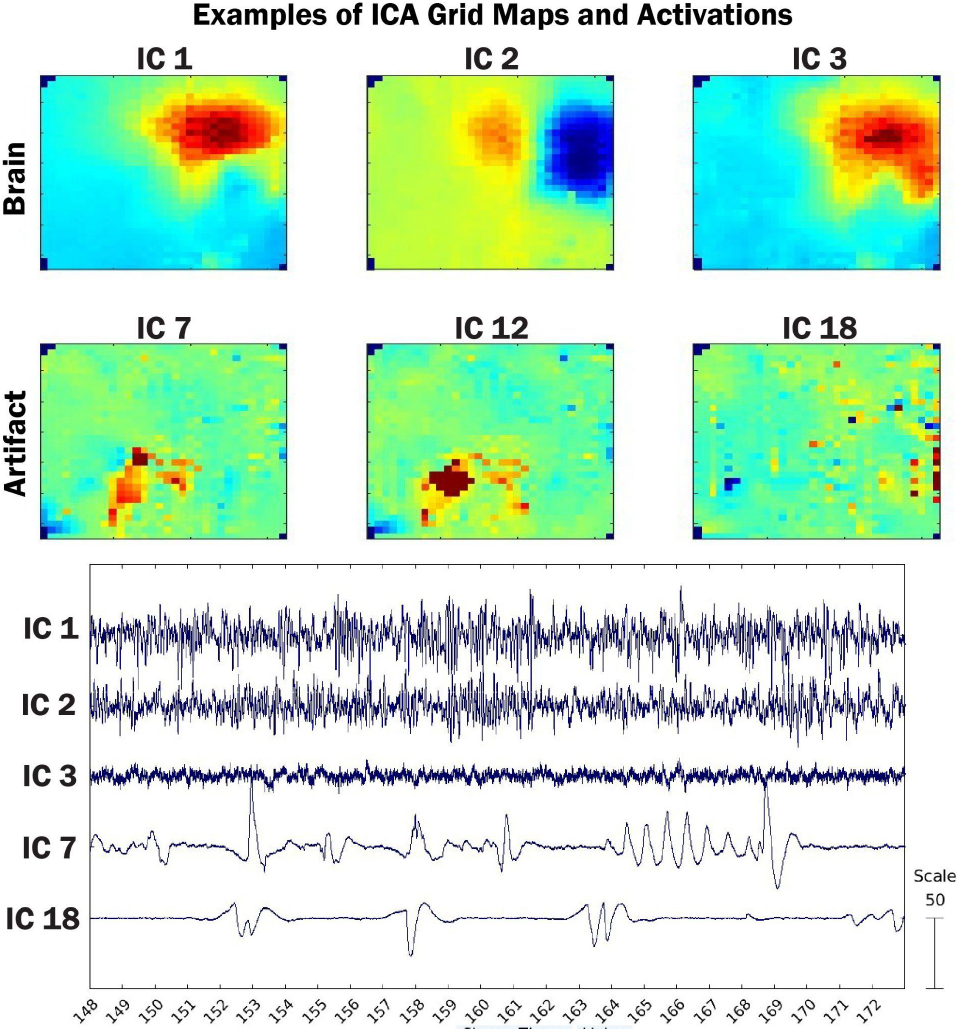
Exemplar inverse weights of Independent Components μECoG grid.

The 582 (of 807) ICs appearing most compatible with cortical activity were retained for use in the simulation experiment. On average, these accounted for 44.6% of the variance of the whole (807-channel) data after baseline correction. Figure 2 shows the percent variance accounted for (pvaf) at each grid channel location.

**Figure 2:**
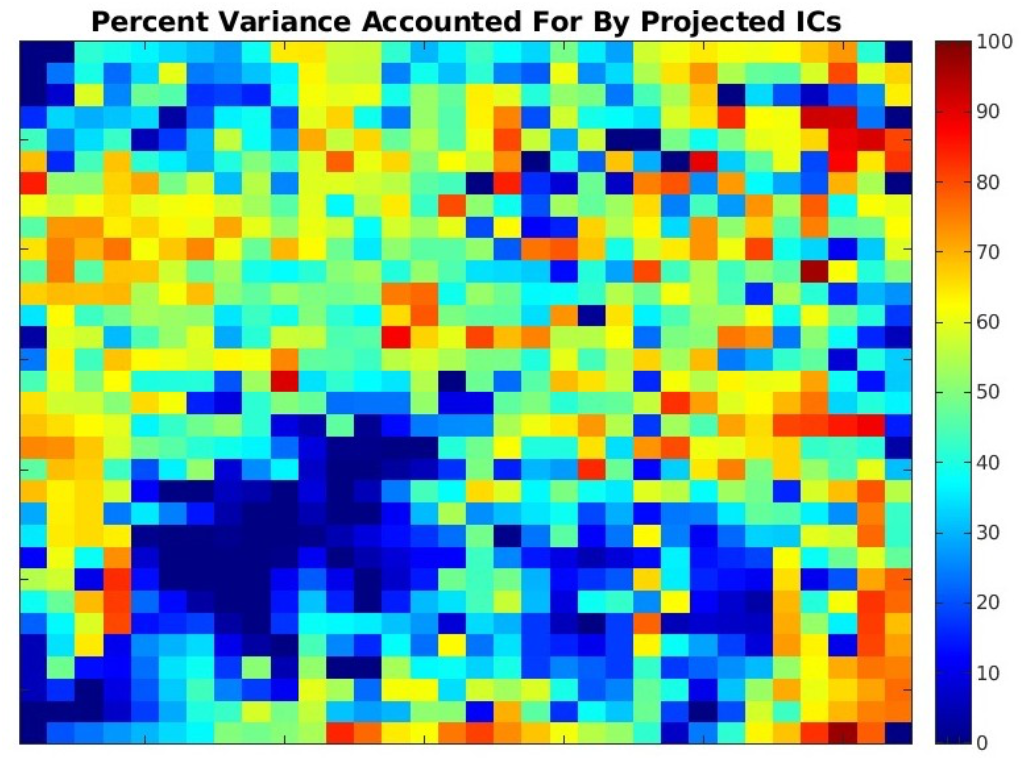
Percent recorded μECoG variance accounted for by the 582 ICs retained after artifact-IC rejections.

### C. EEG Simulation

The retained 807 μECoG grid channels were then forward-projected to simulated scalp EEG channels to approximate what might be recorded if the μECoG activity active under/near the 3-by-3 cm^2^ patch were the only source of scalp EEG activity. Forward-projection to the scalp EEG space used a high-definition Boundary Element Method (BEM) electrical head model with ~80k source nodes constructed from an actual MR head image (not of the actual study participant using the Neuroelectromagnetic Forward Modeling Toolbox [11]. For simulation, the ECoG grid was placed on the imaged cortex in a location (over left sensorimotor cortex) comparable to that reported in [8]. Each node of the forward model cortical surface mesh represented a potential source current field normal to the cortical surface. The average inter-node distance in our high-definition head model was thereby 1.3 mm, compared to the 8-mm inter-node distance used in many EEG head models (typically having ~4k nodes).

Electrical activity was projected to the nearest source model cortical surface nodes, with projection strength inversely proportional to electrode-node distance. Then node source activity was projected to the EEG electrode space using the Lead Field Matrix (LFM) for this head model. The LFM accounts for brain, cerebrospinal fluid, skull shape and electrical conductance properties. Using this LFM, we projected the cortical activities to 207 simulated EEG channels.

### D. EEG analysis

We quantified EEG activity range, standard deviation, and data-length-normalized Shannon entropy for each channel. Here, we used Shanon entropy and entropy interchangeably. We z-scored these metrics to remedy dataset biases. We also identified the average spectral power across channels in delta (1-4 Hz), theta (4-8 Hz), alpha (8-13 Hz), beta (13-30 Hz) and gamma (30-100 Hz) bands. We estimated the Euclidean distance between the centroid of the μECoG patch and each EEG sensor, then built a regression model between observed amplitudes and electrode-patch distance. We assumed the reference for the EEG sensors are located on an electrically inactive surface and far from the EEG sensors themselves. The statistical significance level for regression analysis was p=0.05.

## III. Results

The results show that high-density ECoG grid activity could be decomposed using AMICA into 805 maximally temporally independent source activations with unique grid maps. This initial finding suggests that even in a single 32 × 32 mm patch of cortex, many independent activity patterns are represented. The maximally-independent IC activity patterns sum to dynamically moving activity patterns across the μECoG grid (Figure 1 and Appendix A). There were also ~220 artifactual sources (for example, ICs 7, 12 and 18 in Figure 1) accounting for instrumental or biological artifact activity. Animations of the cortical grid and scalp montage summed and single IC activations (see Appendix) reveal dynamic patterns of activity in various frequency bands. The flow of activity suggests both travelling waves and independent emerging and receding potential spatial patterns.

We retained the subset of ICs (582 out of 805) that appeared to have a cortical origin to backproject first to the μECoG grid (using the pseudoinverse transform) and then to the scalp electrodes. This maximized chances that the μECoG projections to the simulated scalp electrodes through the human forward-problem head model accurately simulated signal loss and spread through head tissues (brain, CSF, skull, scalp). The resulting scalp signals simulated data recorded from a small patch of cortex propagated to the full scalp surface.

Our results show how each (3×3 cm) patch of cortex projects activity to all electrodes of a scalp montage, and therefore to all scalp channels derived from that montage (Figure 3). Notably, here the simulated scalp EEG signal metrics showed strongest projections not in the scalp area directly over the μECoG patch (over the left sensorimotor cortex), but rather scalp signals recorded from the left parietal and right frontal areas.

**Figure 3:**
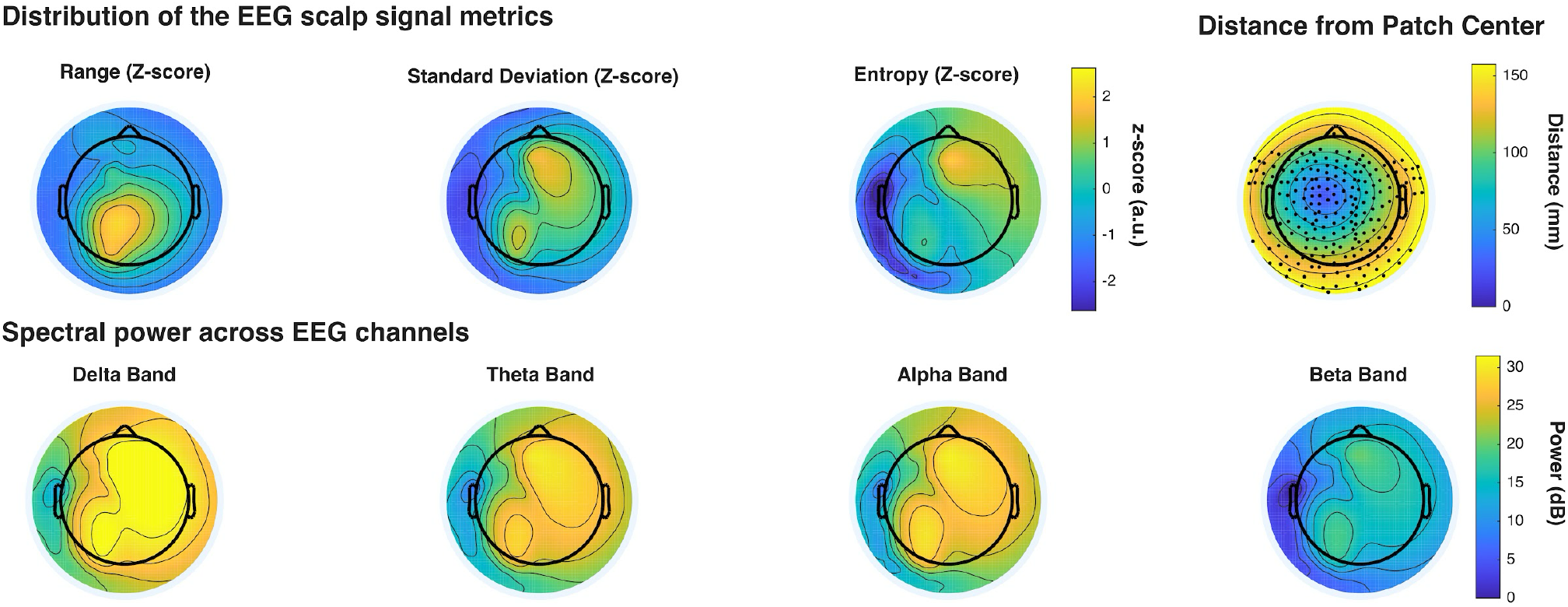
**Top row:** Topographic plots showing topographies of normalized Range, Standard Deviation and Entropy of the simulated scalp signals projected from the uECoG recording. Positions of the simulated scalp electrodes are shown on the upper right scalp map. In these maps the circular head boundary (dark circle) represents the nasion-ear canal-inion head plane. **Bottom row:** Maps of mean log spectral power in the scalp-projected data.

Similarly, spectral power distributions in all frequency bands were strongest in left parietal and right frontal scalp areas (Figure 3, lower row), likely resulting from tangentially flowing activity originating in the central sulcus beneath the grid (captured in part by ICs 1 and 3 of Fig. 1). IC numbers represent respective ranking by variance expressed. Thus, these were two of the three ICs with strongest scalp projections.

Similarly, scalp EEG signal metrics (signal range, standard deviation, and entropy) did not decrease monotonically with distance of the EEG sensors from the μECoG patch (Figure 4). Grouping the EEG sensors into 10 bins based on their distances from the center of the μECoG patch, only signal range (max-min, z-scored) had a decreasing trend with scalp electrode distance from the patch (Figure 4).

**Figure 4:**
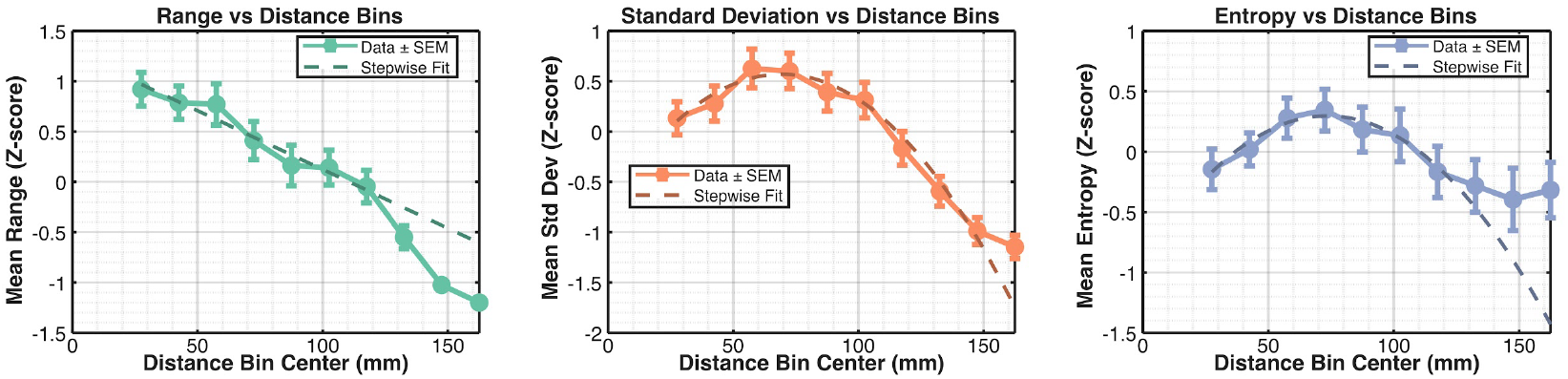
The scalp EEG signal metrics grouped into 10 bins versus the bin distance revleas non-linear relation between the scalp EEG metrics and the EEG electrode distance to the μECoG patch center.

Using MATLAB’s stepwiselm function to determine the trend between the binned signal metrics and the distance of the bin center to the μECoG patch showed, correspondingly, that EEG signal range was linearly correlated with electrode-to-grid distance, while signal standard deviation and entropy showed an inverse quadratic trend.

Regression analyses of the signal metric for individual EEG channels confirmed the overall inverse relation between the EEG signal metrics and the distance of the EEG sensors to the μECoG grid center. However, the degree (or slope) of this relation was considerably different across range, standard deviation, and entropy measures (Figure 5). Interestingly, the Log power spectral density for the delta, theta, alpha, beta, and gamma frequency bands demonstrated a similar degree (slope) of decay power decay with the increase of distance between the EEG electrode and μECoG patch, as depicted by the parallel regression lines for the different frequency bands despite that each regression line had a different intercept.

**Figure 5:**
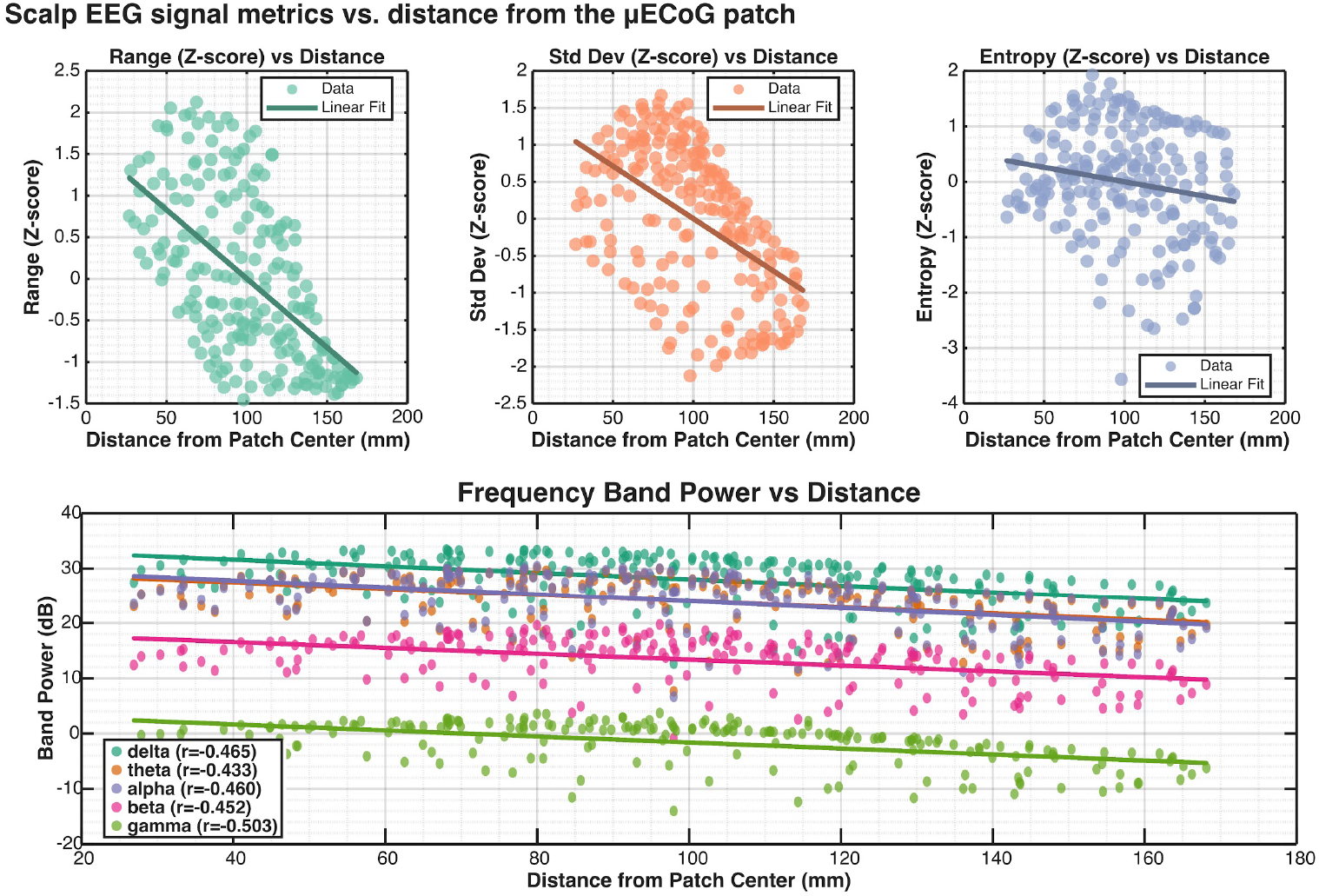
The relation between the EEG signal metrics **(top panel)** and spectral power **(bottom panel)** to the EEG electrode distance from μECoG patch center.

## VI. Discussion

We successfully showed that the μECoG data can be readily decomposed into independent components distinguishing the artifactual and biological sources. While confirming the number of maximally independent sources within the ~580 non-artificial sources require further analysis of the sources using pairwise mutual information (PMI) between the ICs, the dynamic movement of μECoG grid activity suggests that the underlying activity might not be the result of “static” LFPs. The projection of the μECoG activity through the forward head model to the scalp EEG channel space supported our hypothesis that the cortical activity is recorded far beyond the closest EEG electrodes.

LFPs are often assumed and modeled as stationary current dipoles with specific waveform characteristics. Our preliminary results challenge this notion as the recorded μECoG data as well as the decomposed sources do not resemble stationary, but rather moving across the grid. The dynamic nature of these sources suggests that conventional ICA analysis, which assumes temporal stationarity, may not fully capture the complex spatiotemporal dynamics observed in cortical activity. Sources that exhibit non-stationary behavior, such as traveling waves or dynamically shifting activation patterns, may be inadequately decomposed or potentially missed entirely by standard ICA approaches. Deconvolutive ICA methods, which can account for temporal delays and non-stationary mixing processes, may provide superior decomposition of such dynamic cortical activity patterns by relaxing the instantaneous mixing assumption inherent in conventional ICA.

The results from the simulated EEG showed that the μECoG activity can be detected in very far EEG electrodes as well as the close-range EEG electrodes. This finding challenges the notion of attributing the EEG channel activity to the closest cortical area, which is unfortunately a common method in EEG research. However, several signal processing approaches can address this volume conduction issue to varying degrees. Spatial filtering techniques such as the Laplacian filters (Hjorth 1979) can enhance local signal characteristics by computing spatial derivatives that reduce the influence of distant sources, though they cannot completely eliminate volume conduction effects. More sophisticated approaches including ICA-based source separation and dipole-based source estimation methods can more effectively disentangle the contributions of multiple cortical areas to scalp recordings, providing more robust separation of spatially distributed neural activities. Importantly, while the widespread projection of cortical activity poses challenges for localizing brain sources, it also represents an advantage: each EEG channel contains information from a broad range of cortical areas, potentially allowing detection of neural activity from regions that might otherwise be considered too distant to influence scalp recordings.

Limitations of this study include several methodological considerations that, while present, do not fundamentally compromise our main findings. First, we did not use a subject-specific forward head model, as MRI data for the actual participant was not available. However, the high-definition head model employed (with ~80k nodes versus typical ~4k node models) provides sufficient anatomical detail to accurately model volume conduction effects, and the specific head geometry differences would primarily affect absolute amplitude scaling rather than the relative projection patterns that are central to our conclusions. Second, the projection of μECoG electrode activity to the lead field matrix nodes only accounted for the Euclidean distances between electrodes and cortical surface nodes, not the angle between the electrode surface and cortical normal vectors. This simplification may introduce minor errors in projection strength estimation, but given the 1-mm inter-electrode spacing and the relatively flat cortical surface under the grid, angular variations are minimal and unlikely to substantially alter the overall projection patterns observed. Third, our analysis was limited to data from a single subject and recording session, which restricts generalizability. However, the biophysical principles of volume conduction that underlie our findings are universal, and the observed projection patterns are consistent with established principles of electromagnetic field propagation through head tissues.

Future directions include more comprehensive analysis of the μECoG independent components using advanced ICA post-processing techniques such as pairwise mutual information (PMI) analysis and mutual information reduction (MIR) to better understand the dimensionality and independence of cortical source subspaces. These methods can provide quantitative measures of true source independence and help identify potentially over-decomposed or residually dependent components. Additionally, implementing deconvolutive ICA approaches could help disentangle temporally dynamic cortical processes and partially relax the stationarity assumptions inherent in conventional ICA, potentially revealing traveling waves and other non-stationary cortical phenomena that may be obscured by instantaneous mixing models. A particularly valuable extension would involve EEG simulation using multiple active μECoG patches distributed across different cortical regions, followed by comprehensive source estimation analysis of the resulting simulated scalp recordings. Such multi-patch simulations would provide insights into how well current source localization methods can recover known cortical activity patterns in the presence of realistic multi-source interference and would offer a rigorous validation framework for EEG source estimation techniques under controlled conditions where ground truth cortical activity is precisely known.

## V. Conclusion

This study demonstrates that cortical electrical activity recorded from a small (3×3 cm) sensorimotor patch projects broadly across the scalp surface, challenging conventional assumptions about the spatial specificity of EEG channel recordings. Using μECoG data as ground truth cortical activity and forward-projecting through a high-definition head model, we showed that scalp EEG signals reflect complex mixtures of distributed cortical sources rather than primarily local activity beneath recording electrodes. The 582 independent components derived from the μECoG array revealed dynamic, non-stationary cortical activity patterns that projected most strongly to parietal and frontal scalp regions rather than directly overhead, illustrating how volume conduction creates counterintuitive projection patterns. These findings underscore the critical importance of source-level analysis methods for accurate interpretation of EEG data and suggest that channel-level approaches may fundamentally misattribute the cortical origins of observed electrical activity. Our results provide a neurophysiological foundation for improving EEG-based diagnostic and research applications by emphasizing the distributed nature of cortical-to-scalp signal propagation and the necessity of advanced signal processing methods that account for volume conduction effects.

## Acknowledgment

We thank Prof. Shadi Dayeh of UCSD Engineering for permission to use these data for further research.

## Notes

* Research supported by the Swartz Foundation

### Competing Interest Statement

The authors have declared no competing interest.

